# Longitudinal Automatic Segmentation of Hippocampal Subfields (LASHiS) using Multi-Contrast MRI

**DOI:** 10.1101/759217

**Authors:** Thomas Shaw, Ashley York, Maryam Ziaei, Markus Barth, Steffen Bollmann, for the Alzheimer’s Disease Neuroimaging Initiative

## Abstract

The volumetric and morphometric examination of hippocampus formation subfields in a longitudinal manner using *in vivo* MRI could lead to more sensitive biomarkers for neuropsychiatric disorders and diseases including Alzheimer’s disease, as the anatomical subregions are functionally specialised. Longitudinal processing allows for increased sensitivity due to reduced confounds of inter-subject variability and higher effect-sensitivity than cross-sectional designs. We examined the performance of a new longitudinal pipeline (Longitudinal Automatic Segmentation of Hippocampus Subfields [LASHiS]) against three freely available, published approaches. LASHiS automatically segments hippocampus formation subfields by propagating labels from cross-sectionally labelled time point scans using joint-label fusion to a non-linearly realigned ‘single subject template’, where image segmentation occurs free of bias to any individual time point. Our pipeline measures tissue characteristics available in *in vivo* high-resolution MRI scans, at both clinical (3 Tesla) and ultra-high field strength (7 Tesla) and differs from previous longitudinal segmentation pipelines in that it leverages multi-contrast information in the segmentation process. LASHiS produces robust and reliable automatic multi-contrast segmentations of hippocampus formation subfields, as measured by higher volume similarity coefficients and Dice coefficients for test-retest reliability and robust longitudinal Bayesian Linear Mixed Effects results at 7 T, while showing sound results at 3 T. All code for this project including the automatic pipeline is available at https://github.com/CAIsr/LASHiS

## 1. Introduction

The hippocampus formation is a brain structure generating large interest and research activity due to its implication in memory, psychiatric and neurological disorders including Alzheimer’s Disease (AD; Daulatzai, 2013; Fotuhi et al., 2012), Motor Neurone Disease (Machts et al., 2018) and depression (Sapolsky, 2001), and especially its functional and structural changes in ageing (Fraser et al., 2015). Due to the hippocampal formation’s vulnerability in neurodegenerative disease, and its involvement in neurogenesis (specifically within the dentate gyrus [DG], Erickson et al., 2011), precise volumetric and morphometric measurements of hippocampus formation are important for clinical studies and ageing research. Recent work has focussed on the hippocampus formation subfields, which are impacted differentially in neurodegeneration and disease (e.g., Machts et al., 2018). Volumetric and morphometric examination of these hippocampus subfields, especially in longitudinal studies, may lead to more sensitive biomarkers of disorder and the progress of the diseases (Adler et al., 2018; Boutet et al., 2014; Henry et al., 2011; Kerchner et al., 2012; La Joie et al., 2013; Maruszak & Thuret, 2014; Pluta et al., 2012).

Hippocampus subfields are functionally and cytoarchitectonically disparate (Andersen, 2007; Daulatzai, 2013; de Flores et al., 2019; Fotuhi et al., 2012) with heterogeneous cellular composition. The four Cornu Ammonis (CA) subfields each have regional variations in pyramidal cells, creating structural differences between these subfields, which can be reflected to a degree in differing contrast and intensity signals in magnetic resonance imaging (MRI) scans with sufficient sensitivity and spatial resolution, i.e., at high enough field strengths (3 Tesla [T] and above; e.g., Duvernoy et al., 2013; Naidich et al., 2003). As distinct cellular differences between subfields are only observable *ex vivo* and translate into only subtle differences in the MR signal, it is difficult to characterize these tissue classes at lower field strengths and routine MRI sequences due to low signal to noise ratio (SNR) and imaging artefacts (D. Wang & Doddrell, 2005).

Following from the above challenges, changes in small brain structures have been successfully studied at ultra-high field (UHF) using MRI sequences with different contrasts (multi-contrast MRI) and allowed remarkable details for imaging *in vivo* (Fracasso et al., 2016). UHF MRI enables the increased spatial resolution necessary to characterize tissue differences *in vivo*, and in reasonable acquisition times. Previous UHF *in vivo* hippocampus subfield segmentation studies (for review, see Giuliano et al., 2017) utilise ‘dedicated’ sequences (e.g., single- or multi-echo Gradient Echo, Turbo-Spin Echo [TSE]) that exhibit varying intensity and contrast characteristics for different tissue classes due to multiple refocusing pulses. Consequently, the subfields of the hippocampus are observable in these dedicated sequence types (Marques & Norris, 2018; Winterburn et al., 2013).

Advances in MRI acquisition techniques and image analysis methods have made automatic segmentation of hippocampus subfields possible. More recently, fully automatic hippocampus subfield pipelines including Freesurfer’s hippocampus subfields method (Iglesias et al., 2015) and Automatic Segmentation of Hippocampus Subfields (ASHS; Yushkevich et al., 2015) have been released as open-source segmentation software that combine several computational methods to achieve more reliable and precise results.

While both Freesurfer and ASHS have been applied in various studies (Chiappiniello, 2018; Iglesias et al., 2016; Pluta et al., 2012; Yushkevich et al., 2015), generally, segmentation errors cannot be avoided in practice. The cross-sectional variant of Freesurfer accounts for contrast differences in input images while leveraging a combination of T1w and T2w contrasts for defining hippocampus segmentation. The underlying assumption of the Freesurfer scheme is that the spatial distribution of brain structures will be consistent with the *in vivo* and *ex vivo* data in the atlas package, and spatial distributions of brain structures are homogenous within all scanned populations. A longitudinal variant of the hippocampus subfield method from Freesurfer (Iglesias et al., 2016) has also been introduced, which decreases residual (within-subjects) variability by allowing each participant to act as their own control. However, this method does not incorporate T2w information for labelling. It has been shown previously that T1w information generally does not contain signal that differentiates hippocampus subfields (Winterburn et al., 2013), including - in T2w contrast - the hypointense band of cells that separates the dentate gyrus (DG) from the CA regions known as the *stratum radiatum lacunosum moleculare*.

Longitudinal processing allows for increased sensitivity (Fitzmaurice et al., 2011) due to reduced confounds of inter-subject variability and higher effect-sensitivity than cross-sectional designs. In image processing pipelines, longitudinal processing avoids many issues of secular trends inherent to cross-sectional designs, as participants act as their own control. These designs often exploit the knowledge that within-subject anatomical changes over time are usually significantly smaller than changes on an inter-subject morphological scale (Reuter et al., 2012). Longitudinal designs have been used to successfully characterise changes in brain morphometry over time with greater accuracy than their cross-sectional counterparts (Reuter et al., 2012; Reuter & Fischl, 2011; Tustison et al., 2017). These designs avoid many types of image processing bias by transforming images into an intermediate space between time points where interpolation-related blurring is consistent across the time points.

Currently, using ASHS to measure volumes of hippocampus subfields in a single participant at multiple time points does not account for the inherent variability present in cross-sectional methods. The Freesurfer longitudinal hippocampus subfields pipeline is the only dedicated longitudinal pipeline for measuring the volume of hippocampus subfield automatically. However, this method does not utilise the signal and tissue information available with multi-contrast MRI, and in particular, the ‘dedicated’ T2w scan commonly used in measuring the subfields of the hippocampus. We aimed to develop a longitudinal automatic hippocampus subfield segmentation pipeline that incorporates multi-contrast information while being robust to computational errors inherent to purely cross-sectional methods. We then examined the performance of our new longitudinal pipeline (Longitudinal Automatic Segmentation of Hippocampus Subfields [LASHiS]) against three published approaches viz; cross-sectional (FS Xs) and longitudinal (FS Long) Freesurfer hippocampal subfields (V6.0 Dev20181125; Iglesias et al., 2016), and ASHS cross-sectional (ASHS Xs; Yushkevich et al., 2015)

We developed an open-source multi-contrast pipeline that shares commonalities with existing pipelines but can capture multi-contrast information from MRI scans automatically, while avoiding errors common to cross-sectional processing. We integrate several open-source software packages and programs to construct LASHiS and propose the usage of multi-atlas fusion techniques to bootstrap automatic segmentation performance. Our pipeline is implemented with existing tools available through ANTs (ANTs Version: 2.2.0.dev116-gabc03; http://stnava.github.io/ANTs/; Avants,Tustison,&Song,2010)andASHS (https://sites.google.com/site/hipposubfields/; Yushkevich et al., 2015). Our pipeline and all associated code can be found at https://github.com/CAIsr/LASHiS.

## 2. Methods and materials

### 2.1 Towards Optimising MRI ChAracterisation of Tissue (TOMCAT) imaging data

Seven healthy participants (age: M = 26.29, SD = 3.35, sex: 3 female, 4 male) were scanned using a 7 T whole-body research scanner (Siemens Healthcare, Erlangen, Germany), with maximum gradient strength of 70 mT/m and a slew rate of 200 mT/m/s and a 7 T Tx/32 channel Rx head array (Nova Medical, Wilmington, MA, USA) in three sessions with three years between session one and two, and 45 minutes between two and three, allowing for a scan-rescan condition. The study was approved by the university human ethics committee and written informed consent was obtained from the participants. Participants were scanned using a 2D TSE sequence (Siemens WIP tse_UHF_WIP729C, variant: tse2d1_9), TR: 10300 ms, TE: 102 ms, FA: 132°, FoV: 220 mm, voxel size of 0.4 × 0.4 × 0.8 mm^3^ Turbo factor of 9; iPAT (GRAPPA) factor 2, acquisition time (TA) 4 minutes 12 seconds. The scan was repeated thrice over a slab aligned orthogonally to the hippocampus formation. An anatomical whole-brain T1w scan using a prototype MP2RAGE sequence (WIP 900; Marques et al., 2010; O’Brien et al., 2014) at 0.75 mm isotropic voxel size was also acquired (TR/TE/TIs = 4300 ms / 2.5 ms / 840 ms, 2370 ms, TA = 6:54). At the first time point, the data was acquired as part of a larger study (Bollmann et al., 2018), and the nominal resolution was 0.5 mm isotropic with the same parameters. For all subsequent processing, all MP2RAGE images for the first time point were resampled to 0.75 mm isotropic using b-spline interpolation. TSE images were resampled to 0.3 mm isotropic and motion-corrected using non-linear realignment (Shaw et al., 2019) to ensure all segmentation strategies had an equivalent chance of succeeding. We have previously explored the effects of non-linear realignment on hippocampus subfield segmentation (Shaw et al., 2019) and have found the method to be beneficial for segmentation reliability and sharpness.

Non-linear realignment is a process of minimum deformation averaging of multiple repetitions of the same sequence, which boosts SNR and image sharpness. This method works best when input images are isotropic (in this case interpolated using 3^rd^ order b-splines). Inter-scan movement before the initial interpolation could lead to unwanted artefacts further downstream in the realignment process. However, non-linear realignment has been shown to iteratively converge more readily to ‘crisp’ isotropic voxels with distinct features, and the realignment procedure ensures that only anatomically consistent features of the three TSE images are retained, which motivated our choice.

#### 2.1.2 ADNI-3 MRI data

As a follow up study for the small *n* TOMCAT dataset, we used longitudinal data from the Alzheimer’s Disease Neuroimaging Initiative (ADNI)-3 (adni.loni.usc.edu). The ADNI was launched in 2003 as a public-private partnership, led by Principal Investigator Michael W. Weiner, MD. The primary goal of ADNI has been to test whether serial magnetic resonance imaging (MRI), positron emission tomography (PET), other biological markers, and clinical and neuropsychological assessment can be combined to measure the progression of mild cognitive impairment (MCI) and early Alzheimer’s disease (AD).

At 3 T, either an anatomical 1 mm^3^ isotropic full-brain T1w MP-RAGE or Accelerated SPGR sequence was acquired. A high-resolution ‘dedicated hippocampus slab’ T2w TSE or FSE was also acquired at 0.4×0.4×1 mm. Specific details and acquisition parameters are given in http://adni.loni.usc.edu/wp-content/uploads/2017/07/ADNI3-MRI-protocols.pdf. The data was downloaded in October of 2019 and consisted of 112 participants that had both T1w and T2w scans and been scanned two or more times (fulfilling the longitudinal condition). 17 participants had three time points. 79 participants were included in the final analysis, as some data sets could not be processed due to computational errors in one or more of the pipelines or time points, resulting in 11 participants with three time points and 68 with two time points. We note that Freesurfer produced the most failures in processing (n = 79), with LASHiS and ASHS completing 92 cases each. We included all results for each method in the GitHub repository associated with this work. In total, 43 females and 36 males were processed meeting the following diagnostic criteria: 12 late mild cognitive impairment (LMCI), 31 cognitively normal (CN), 8 mild cognitive impairment (MCI), 7 significant memory concern (SMC), and 20 early mild cognitive impairment (EMCI) (Age M = 77.68, SD = 8.76).

Data were converted to BIDS using BIDSCoin (https://github.com/Donders-Institute/bidscoin), and pre-processed identically to the TOMCAT dataset, with the exception of resampling to 0.4 mm isotropic instead of 0.3 mm isotropic for the T2w scan.

### 2.2 Longitudinal Assessment of Hippocampal Subfields (LASHiS)

#### 2.2.1 Atlas Construction

The entire LASHiS pipeline is described in Figure 1. Optionally, the ASHS pipeline can be optimised through the incorporation of a group-specific atlas. Similarly, creation of a group-specific atlas is a benefit to our proposed method. This atlas is comprised of a representative pool of subjects (approximately 20-30 participants), manually labelled, and passed through the ASHS_train pipeline described in Yushkevich et al. (2015). Essentially, the manual segmentations are used as inputs (priors) for the joint-label fusion (JLF) algorithm in subsequent segmentations, and to train classifiers for the ASHS cross-sectional pipeline. Creating a group-specific atlas (of 20-30 subjects) would be beneficial for large longitudinal studies, as segmentation training would be performed on group-specific characteristics. However, having a group specific atlas is generally not necessary for robust performance of ASHS (Xie et al., 2018).

**Figure 1.**
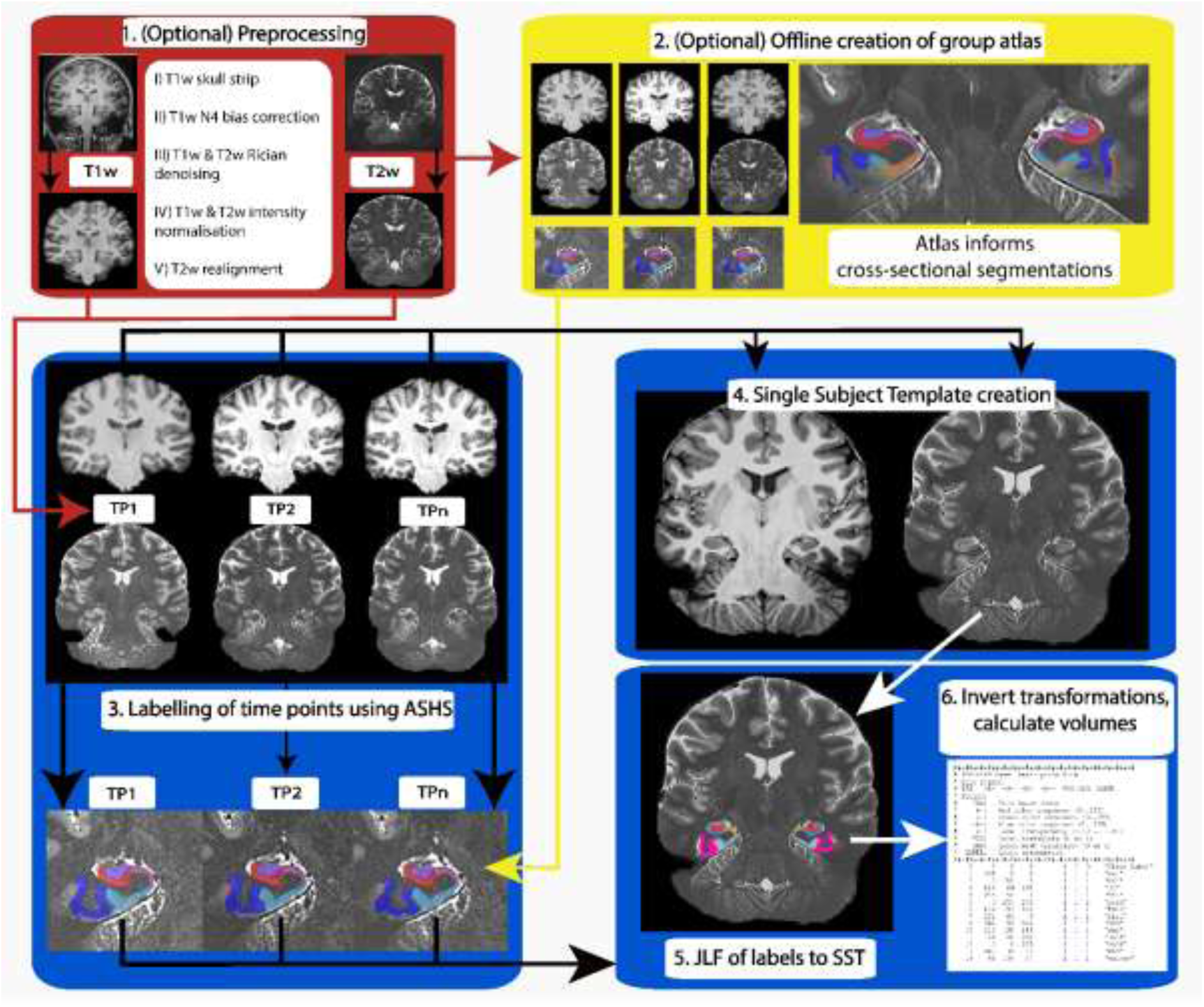
Longitudinal Automatic Segmentation of Hippocampus Subfields (LASHiS) pipeline. The pipeline consists of the following steps with the input of any number of T1w and T2w individual time points per participant: 1) (In red; optional) pre-processing of both T1w and T2w scans. 2) (In yellow) Offline construction of a sample-specific atlas for LASHiS. 3) Labelling of individual time points of each subject using ASHS and a representative atlas (or an atlas created in [2]) to yield hippocampus subfield estimates. 4) Construction of a multi-contrast Single Subject Template (SST). 4) JLF of each of the individual time point labels to the SST using both contrasts and individual hippocampus subfield labels to produce a labelled SST. 5) Application of the inverse subject-to-SST transformations to SST labels. 6) Measurement and calculation of subfield labels in subject-space.

#### 2.2.2 Pre-processing and cross-sectional processing

The ASHS Xs pipeline has been previously proposed and discussed (Yushkevich et al., 2015). Briefly, the pipeline labels hippocampus subfields of a given T1w and dedicated T2w scan covering the hippocampus subfields. This approach leverages a multi-atlas segmentation method and corrective learning techniques to segment (usually 3 T or 7 T) MRI data. The process involves first training existing manually labelled *in vivo* atlases of T2w scans. These trained atlas packages inform labels for new *in vivo* T1w and dedicated T2w scans. ASHS provides many of these atlases at https://www.nitrc.org/projects/ashs. These open source atlases may be replaced with a group-specific atlas as proposed in 2.1.1. The T2w input scan is usually acquired anisotropically with reduced resolution along the major axis of the hippocampus subfield and high in-plane resolution. The spiral structure of the hippocampus formation does not change rapidly along its major axis, which motivates this parameter choice (Iglesias et al., 2016). ASHS Xs employs similarity-weighted voting for learning segmentation priors and JLF for multi-atlas segmentation prior to classification. In the segmentation protocol, weighted voting at the voxel level derives ‘strong’ segmentation choices for the target image (Yushkevich et al., 2015).

For pre-processing of all data, we included modified pre-processing steps based on the ANTs cortical thickness pipeline (Tustison et al., 2014) and our previous work (Shaw et al., 2019). These steps were incorporated to ensure consistent segmentation results across participants and included:

I. Skull stripping (i.e., ROBEX; Iglesias, Liu, Thompson, & Tu, 2011) of the T1w scan for removal of background tissue and artefacts that may result in registration errors further downstream
II. N4 bias correction (ANTS version 2.20.dev116-gabc03; Tustison et al., 2010) of the T1w scan that mitigates low spatial frequency variations in the data
III. Rician denoising of T1w and T2w scans (Manjón et al., 2010), which has been shown to reduce high-frequency Rician noise in MRI scans (Tustison et al., 2017)
IV. Intensity normalisation of T1w and T2w scans to the atlas using histogram matching (Nyúl et al., 2000)
V. If multiple repetitions of the dedicated T2w scan are available, non-linear realignment of these scans to reduce motion artefacts and increase the sharpness of the scans as in Shaw et al. (2019)

LASHiS derives initial segmentations of each time point cross-sectionally using the ASHS pipeline with an atlas package similar to the subjects’ intensity and spatial characteristics. This yields an atlas-defined number of hippocampus subfield labels. Due to our small sample size, it was not possible to create a bespoke atlas for validation. Therefore, we utilized ASHS (V2.0) with the Penn Memory Center 3 T ASHS Atlas (Yushkevich et al., 2015). We motivated this choice based on Xie et al (2018), who found that atlas composition does not significantly affect segmentation between 7 T and 3 T, and the contrast and intensity profiles of the scans in the 3 T atlas are similar to the TSE scans we collected in the present study.

#### 2.2.3 Single-subject template (SST)

In parallel to cross-sectional processing, a minimum deformation average multi-contrast template of average intensity and shape is created in accordance with Avants et al. (2010). This template serves as an intermediate between any *n* time points of a subject and is biased equally to any given time point. All subsequent processing of hippocampus volumes is done in the space of the SST in order to treat all time points in the same way. We have also found previously that combining scans in this way increases segmentation consistency and image sharpness (Shaw et al., 2019).

#### 2.2.4. Joint-Label Fusion (JLF) and longitudinal estimations of segmentations

Following SST creation and labelling, and cross-sectional labelling, individual time point multi-contrast scans and their cross-sectionally defined segmentation labels then act as multi-contrast atlases to compute SST labels using JLF. The intended usage of JLF is to propagate manually derived labels to a target image. However, we used JLF with the atlases being *automatically* labelled. We also include the automatically labelled SST as an extra input to increase the power of the method. JLF assigns the spatially varying atlas (input) weights to the SST in a way that accounts for error correlations (Wu et al., 2017) between every *n* pairs of atlases. In this way, no single time point is biased towards the segmentation of the SST, and the SST is labelled based on a weighted vote of the segmentations from each time point and the SST. In our scheme, a working region of interest (ROI) is defined roughly around the hippocampus, non-linearly warped to the space of the SST ROI, and JLF applied with parameters chosen based on (Wang et al., 2013). The inverse of these non-linear transformations is later used for labelling the input images. This approach, therefore, bootstraps cross-sectional segmentations of hippocampus subfields to the SST, and the best fitting labels are chosen based on the intensity and shape characteristics of the SST, not the individual time points.

Subsequently, the inverse of the time point to SST transformations from the JLF piecewise registration is applied to the generated SST hippocampus subfield labels, warping the SST labels to each individual time point. Provided the time point-to-SST registrations are accurate (Avants et al., 2011) and invertible, the reverse normalisation of the labels can be considered a robust and reliable method for transforming the labels to the space of the subject’s time point hippocampus subfield labels. Finally, we used Convert3D (Yushkevich et al., 2006) to measure the new subfield volumes in time point space.

### 2.3 Statistical Evaluation

#### 2.3.1 Hippocampus subfield segmentation methods

We compared the performance of our LASHiS strategy with three other established strategies, and one other exploratory strategy detailed below, examples of the output of each segmentation strategy are given in Figure 2:

1. Cross-sectional ASHS (ASHS Xs), the segmentation strategy described in Yushkevich (2015), was used to compute segmentations for each time point independently in a cross-sectional manner. We utilised segmentation results that incorporated the high-resolution T2w scan information. We modified certain parameters in ASHS to account for our 7 T high-resolution and pre-processed (isotropic) inputs to account for resolution and image size. We used the Penn Memory Center atlas https://www.nitrc.org/frs/?group_id=370 for segmentation due to input-atlas contrast similarities and an increased number of subfield label outputs compared to the available 7 T atlases.
2. Cross-sectional Freesurfer hippocampus subfield segmentation (FS Xs): the method described in Iglesias et al. (2015) was used to compute segmentations for each time point independently in a cross-sectional manner. Due to skull strip failures in recon-all and mri_watershed, the brain mask was replaced with the brain mask created in the pre-processing steps using ROBEX (Iglesias et al., 2011) in order to give Freesurfer the best chance of succeeding.
3. Freesurfer longitudinal hippocampal subfields (FS Long): This pipeline, described by Iglesias et al. (Iglesias et al., 2016) utilises intensity and contrast information from an *ex vivo* manually traced atlas of hippocampal subfields to delineate *in vivo* subfield information. The *ex vivo* atlas is supplemented by an *in vivo* Freesurfer atlas (as described in Kennedy et al., 1989), which informs segmentation of the geometric priors surrounding the hippocampus. In this way, the generative model that classifies hippocampal subfields *in vivo* is calculated from the spatial distribution of the hippocampus and its surrounding brain regions as described in the atlas priors.
4. JLF-free LASHiS (Diet LASHiS): This method is similar to LASHiS, though does not utilise the JLF bootstrapping step or cross-sectional processing, thus reducing processing time by approximately 20%. Instead, the SST is created, labelled using ASHS and a representative atlas package, and labels were reverse-normalised to time point space using the transformations calculated in the template building procedure, as distinct from the SST transformations derived in the JLF step. We incorporated this method to determine the relative importance of the JLF bootstrapping step in our pipeline. This would be considered a standard reverse normalisation segmentation pipeline.

**Figure 2.**
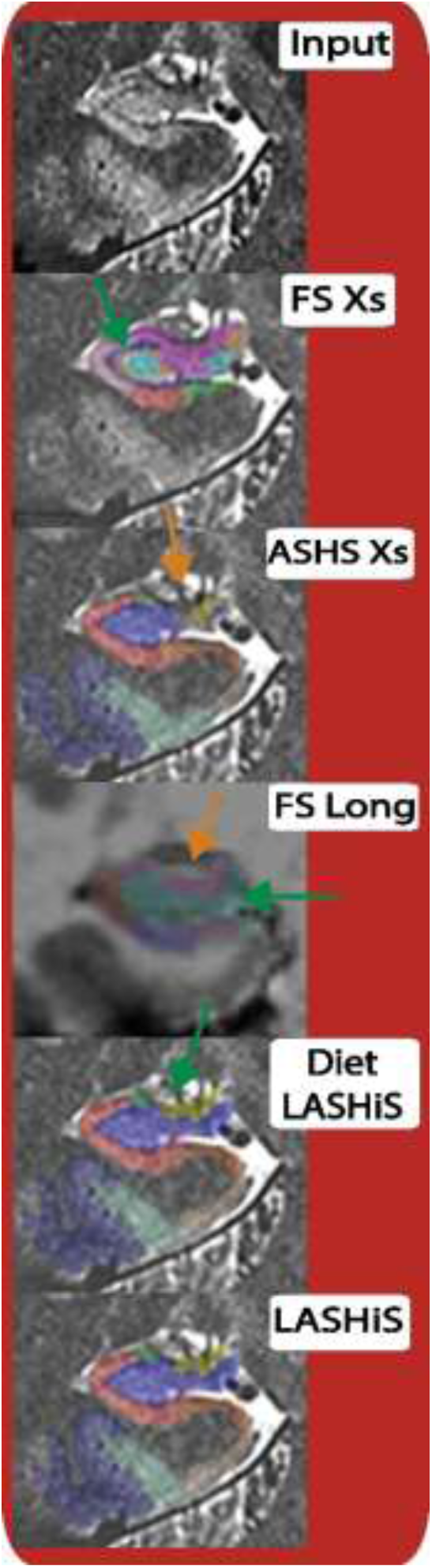
Hippocampus subfield segmentation results (coloured) for a single representative subject for the five tested methods at the same slice in a coronal view. Each segmentation result is overlaid on the high-resolution T2w scan except for FS Long, which utilises a T1w scan for segmentation. Green arrows denote a possible under-segmentation, orange a possible over-segmentation.

#### 2.3.2 Evaluation methods

Here, we evaluate our strategy in line with previously published methods in order to quantify reliability, reproducibility and precision. We reproduced analyses employed by both the FS Long hippocampus segmentation strategy (i.e., test-retest reliability) and longitudinal Bayesian Linear Mixed Effects (LME) modelling (Sorensen et al., 2016).

#### 2.3.3 Experiment one: Test-retest reliability

We evaluated the test-retest reliability of all methods through testing differences between the second and third time point in the TOMCAT dataset. For each participant, we segmented each scan-rescan session with the five segmentation methods and assessed performance based on two metrics: 1) absolute differences in volume estimates for each hippocampus subfield label between scan-rescan acquisitions, and 2) the Sørensen-Dice similarity coefficient (Dice, 1945). We first measured the volume similarity coefficient, which does not rely on segmentation locations (Taha & Hanbury, 2015). This metric does not implicitly rely on overlaps in segmentations (such as Dice overlaps, which can be difficult to measure without bias when comparing between analysis strategies, as in Iglesias et al., [2016]). For completeness, and to have a direct comparison with Iglesias et al. (2016), we also assessed Dice overlaps between time point two and three in all pipelines. The Sørensen-Dice similarity coefficient between two binary masks is described as “twice the number of elements common to both segmentations divided by the sum of the elements in the segmentations”, and is defined as:

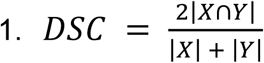

where a perfect overlap between two segmentations (*X* and *Y*) is 1, and no overlap is 0 (Taha & Hanbury, 2015). In LASHiS, Diet LASHiS, FS Xs, and ASHS Xs, the final result of hippocampus subfield labels occurs in a native (input) space. We resampled all labels in these four pipelines to an intermediate space (SST space) using a rigid linear realignment, and calculated Dice overlaps in these cases with the fuzzy Dice counterpart using the freely available EvaluateSegmentation tool (Taha & Hanbury, 2015). There is a bias towards FS Long for having superior Dice overlap evaluation due to the extra interpolation required in these linear realignments, which are not necessary in FS Long. We discuss the implications of this in section 4.1.

We leveraged Bayesian paired *t*-tests in accordance with Rouder, Speckman, Sun, Morey, and Iverson (2009) to assess the differences in subfield changes across the second and third time point using Jamovi, R, and the BayesFactor plugin (Morey & Rouder, 2019; R Core Team, 2019; The Jamovi Project, 2019). In our analyses, BF_10_ > 3 was taken as substantial evidence for the alternative hypothesis, with BF_10_ > 10 taken as strong evidence, and BF_10_ greater than 100 were considered decisive. BF_10_ values between 1 and 3 were considered anecdotal evidence for the alternative hypothesis. In contrast, BF_10_ < 0.33 (or BF_01_ > 3) was considered as substantial evidence for the null, with BF_10_ between 0.33 and 1 providing anecdotal evidence for the null hypothesis in accordance with Lee and Wagenmakers (2013). BF_10_ values can be interpreted to mean that these data are *x* many times more likely to be observed under the alternative hypothesis than the null hypothesis, such that BF_10_ = 3 suggests that these data are 3 times more likely under the alternative hypothesis than the null hypothesis. BF_10_ between 0.33 and 1 can be considered as anecdotal evidence for the null, while values around 1 are non-evidential.

#### 2.3.4 Experiment two: Bayesian Longitudinal Linear Mixed Effects Modelling

To assess relationships between cross-sectional and longitudinal results while accounting for subject-specific trends (Tustison et al., 2017), we quantify between, and within (residual) variability of hippocampus subfield volume. In this experiment, we aimed to assess the utility of each pipeline for measuring each hippocampus subfield and detecting potential biomarkers therein. It is possible to quantify the relative performance of cross sectional and longitudinal pipeline variants with Bayesian LME models (Tustison et al., 2017). Intuitively, the best longitudinal method maximises both within-subject reproducibility and between-subjects variability (to distinguish between sub-groups). Maximising the ratio between between-subject variability and residual variability indicates a good performance. A summary measure of this is the variance ratio, which shows the linear relationship between within- and between-subjects variability, which is a useful measure of performance for longitudinal pipelines. Higher variance ratios indicate optimised prediction and credible intervals for the segmentation quality.

Freesurfer and ASHS provide different outputs in terms of subfield names. To overcome difficulties computing variance values relating to non-overlapping regions, we have concatenated several subfields that share commonalities across all pipelines and present these in Table 1. We excluded subfields that did not share any commonalities across pipelines from the analysis.

**Table 1.** Label names for all hippocampus subfields that share similarities between Freesurfer and ASHS pipelines.

| Combined label name for analysis | Freesurfer label names (FS Xs, FS Long) | ASHS label names (ASHS Xs, Diet LASHiS, LASHiS) |
| --- | --- | --- |
| CA1 | CA1-head & CA1-body | CA1 |
| CA2-3 | CA3-head & CA3-body<sup>†</sup> | CA2 & CA3 |
| DG | GC-ML-DG-head, GC-ML-DG-body, CA4 head, & CA4 body | DG<sup>‡</sup> |
| SUB | Presubiculum-head, presubiculum-tail, subiculum head, & subiculum tail | SUB |

<sup>†</sup>Freesurfer combines estimates of CA2&CA3 as label CA3 in their algorithm.
<sup>‡</sup>ASHS combines estimates of DG and CA4 as label DG in their algorithm.

The subfields chosen in our analysis include Cornu Ammonis (CA) region 1 (CA1), CA region 2 and 3 (which was combined in Freesurfer’s method; CA2-3), Subiculum (SUB; comprising presubiculum and subiculum in the Freesurfer pipeline), and dentate gyrus (DG; comprising of CA4 and DG in the ASHS atlas package). These four subfields are measured for all analyses for left and right sides for a total of eight subfields. Note that LASHiS computes as many labels as in the initial atlas package (usually 14 per side). To obtain more internally consistent measures of volume and to focus results on determining what fraction of hippocampal volume is attributed to each subfield rather than fractions of whole brain volume, we normalised all raw volume values by total hippocampus formation volume (e.g., CA + DG + SUB). We examined subfield results for all comparisons but report significant differences only between the most relevant comparisons: namely between LASHiS and FS Long, as these are the two longitudinal pipelines of interest.

## 3.0 Results

### 3.1 Experiment one: test-retest reliability

We conducted a series of Bayesian paired-sample t-tests in order to test absolute volume differences between the second and third time point. Figure 3 shows differences between methods for volume similarity in this test-retest experiment. We found that LASHiS and Diet LASHiS showed significantly higher volume similarity in all subfields than other methods. Specifically, we found LASHiS to have *decisively* higher (BF_10_ > 100) volume similarity coefficients compared to FS Long in all subfields. ASHS Xs also showed high volume similarity in DG subfields compared to the Freesurfer variants, though with high variability; we observed larger variability in the volume similarity in all other methods compared to LASHiS variants (see Supplementary Figure 1 for subfield variance breakdowns). All other comparisons with LASHiS are detailed in Supplementary Figure 1 and 2.

**Figure 3.**
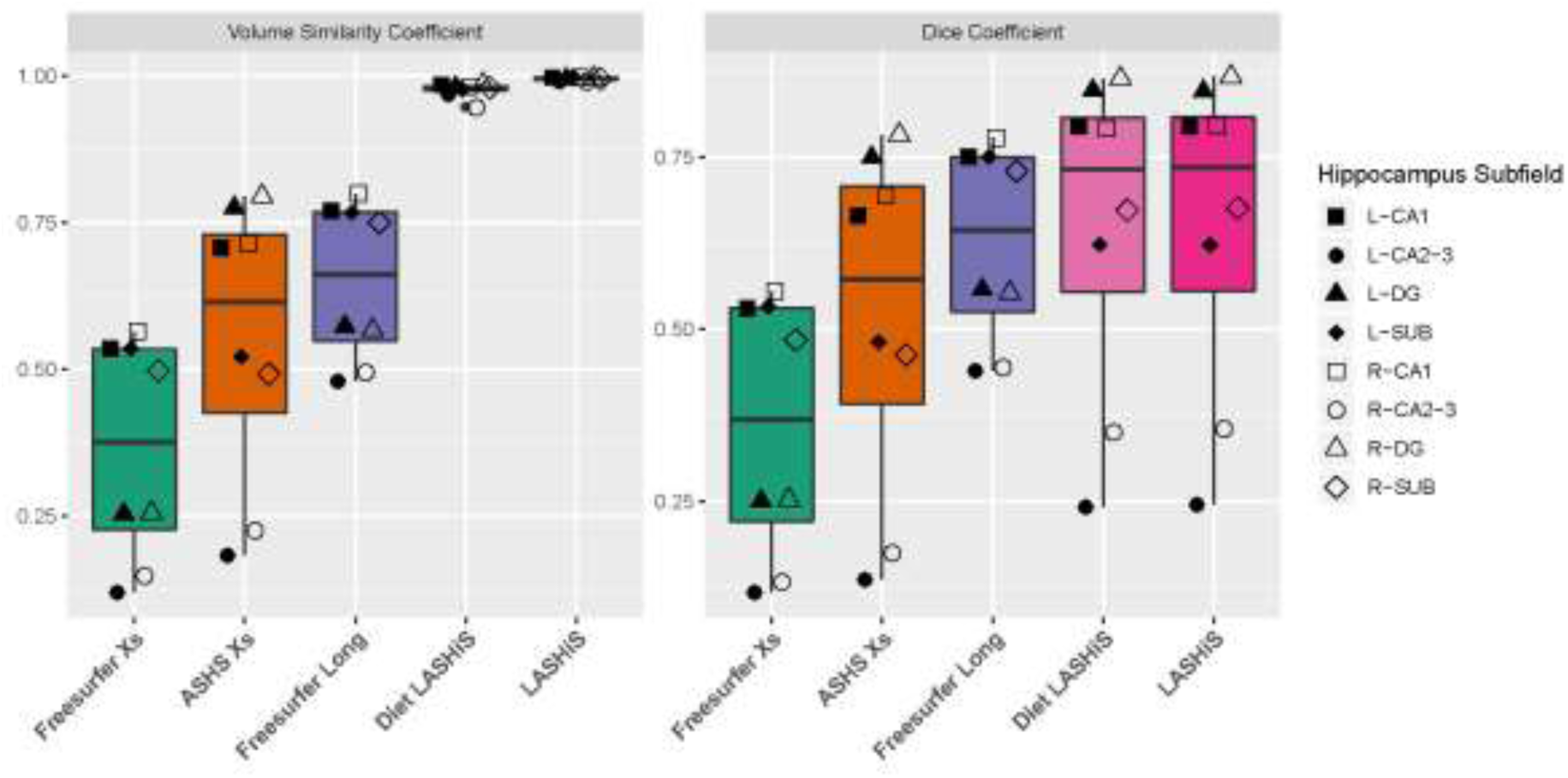
Box plots of volume similarity coefficients (left) and Dice coefficients (right) of each hippocampus subfield for the TOMCAT dataset (7 T). (left, black filled shapes, and right, white filled shapes) from time point two and time point three for each method, where a value of 1 represents perfect overlap between time points, and 0 represents no overlap. Error bars represent overall standard deviation. Freesurfer Xs, ASHS Xs, Diet LASHiS and LASHiS all require resampling to a common space before overlap calculation of Dice scores. Higher scores between time points denote higher subfield overlap between the test-retest conditions.

We next conducted Bayesian paired-sample *t*-tests for Dice overlaps between the segmentation labels in the second and third time point. Figure 3 shows Dice overlap values of each subfield for each method. Note, that Dice scores for LASHiS, Diet LASHiS, Freesurfer Xs and ASHS Xs are negatively affected by the resampling needed to compute the registrations between the two time points, which is not present in the FS Long method. Our results do not fully replicate Iglesias et al. (2016) in terms of mean Dice overlap scores for test-retest reliability, but we found slightly lower Dice overlaps for all subfields in our sample in the Freesurfer methods compared to Iglesias et al. (2016). This discrepancy is potentially due to methodological differences between Iglesias et al. (2016) and our study - scanner hardware (1.5 T rather than our 7 T) and sequence choice (IR-SPGR rather than our MP2RAGE).

In terms of subfield differences, we will detail comparisons only for LASHiS and FS Long, with all method comparisons included in Supplementary Figure 3 and 4. We found that Dice overlaps for LASHiS were higher than FS Long for test-retest reliability *decisively* in Left-DG and Right-DG (BF_10_ > 100) and *anecdotally* in Left-CA1 (BF_10_ > 1). We found no difference between LASHiS and FS Long in Right-CA1, and Right-SUB (BF_10_ < 1). FS Long had *substantially* higher scores than LASHiS for Right-CA2-3, Left-CA2-3, and Left-SUB (BF_10_ > 10).

### 3.2 Experiment Two: Bayesian longitudinal Linear Mixed Effects modelling

We compared the performance of five hippocampus subfield segmentation processing approaches using longitudinal LME modelling to quantify between and residual variability, and the variance ratio of these. Figure 4 provides the 95% credible intervals for the variance ratios in each subfield for each of the pipelines. As noted in Tustison et al. (2017)”, “superior methodologies are designated by larger variance ratios”. For the TOMCAT (7 T) dataset and across subfields, LASHiS has higher variance ratios for Left-CA1, Left-SUB, Right-DG, and Right-SUB. FS Long out-performs LASHiS for Left-CA2-3 and Right-CA1 and ASHS Xs performs best for Right-CA2-3 and Left-DG (followed closely by LASHiS and diet LASHiS). We also note lower values in CA2-3 subfield variance ratios in LASHiS in both hemispheres. For the ADNI dataset (3 T), across subfields we found overlapping 95% credible intervals for all subfields between LASHiS and Freesurfer Long, with the exception of right DG, where LASHiS has significantly higher variance ratios than all other methods including FS Long.

**Figure 4.**
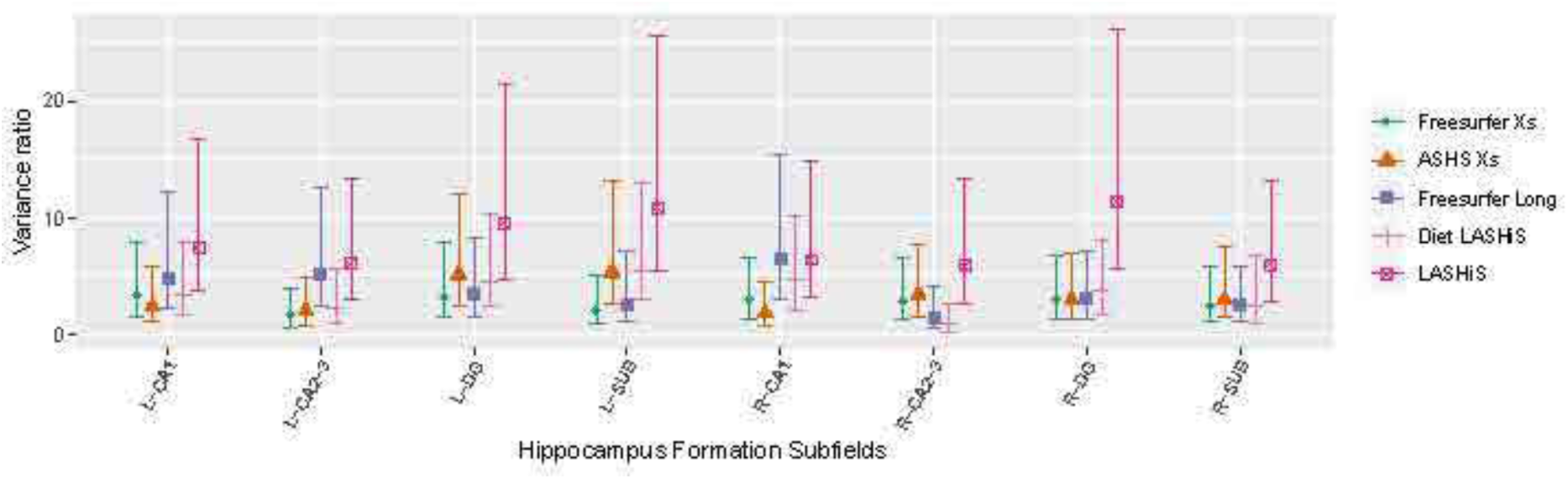
Variance ratio for each subfield (x-axis), for each method (coloured lines) for the TOMCAT dataset (7 T). Values represent the linear regression between residual and between-subjects variability. Therefore, higher values indicate better discrimination between subjects, and higher within-subject reproducibility between the test-retest conditions. Shapes represent the mean variance ratio, with lines denoting the 95% credible intervals for each method.

For the TOMCAT dataset, we found overlapping credible intervals for all pipelines for variance ratios, with obvious trends towards LASHiS as having the highest variance ratios. Figure 4 shows the relative distributions of variance ratios per subfield for each of the assessed pipelines. A trend towards higher variance ratios for LASHiS compared to the other methods can be observed. We also note that variance ratios for Diet LASHiS are comparable to FS Long. There are clear trends towards LASHiS having the superior variance ratios in Left and Right CA1, Left and Right SUB, and Right DG regions. LASHiS has low variance ratio values in Right CA2/3 and therefore has higher variance ratios than all other methods in 5 of 8 subfields. We also provide within and between subject variability for each subfield in Supplementary Figure 5 and 6, respectively, and the overall breakdowns for the variance ratio in Supplementary Figure 7.

For the ADNI dataset, we found that the overall residual variability for Fs Long and LASHiS were generally lower (non-significant) than all other methods (Supplementary Figure 8), resulting in high variance ratios for Fs Long and LASHiS (Figure 5). We note higher between-subjects variance in 6 of 8 subfields in LASHiS compared to FS Long (Supplementary Figure 9). However, all differences were non-significant, except for the variance ratio in right DG (Figure 5), which was significantly higher in LASHiS compared to FS Long. This significant difference was largely driven by the low residual variability in this subfield for LASHiS.

**Figure 5.**
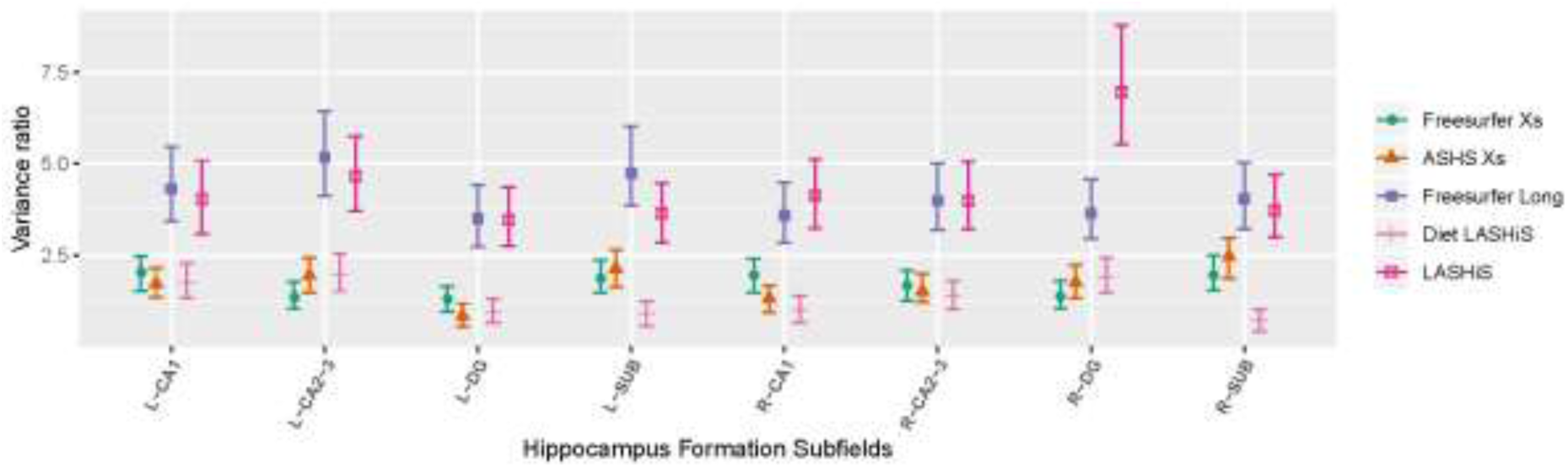
Variance ratio for each subfield (x-axis), for each method (coloured lines) for the ADNI sample (3 T). Values represent the linear regression between residual and between-subjects variability. Therefore, higher values indicate better discrimination between subjects, and higher within-subject reproducibility between the test-retest conditions. Shapes represent the mean variance ratio, with lines denoting the 95% credible intervals for each method.

## 4.0 Discussion

### 4.1 Experiment One, test-retest reliability

The test-retest results highlight the reliability of the LASHiS pipeline. Capitalising on the availability of data from multiple time points to increase SNR in the SST improves the inherent regularisation and prior information for segmentation, as proposed for LASHiS. LASHiS and Diet LASHiS show excellent test-retest reliability for volume similarity coefficients. Deformable registration has been previously used successfully to segment hippocampus structures in groups of participants (Hammers et al., 2007; Hogan et al., 2000). LASHiS benefits from deformable registration-based image segmentation, as the hippocampus is segmented only in SST space. We contrast LASHiS with FS Long, which utilises an SST in order to compute time point segmentations. However, the FS Long SST uses only T1w information, potentially limiting the reliability of the time point to SST registration, and therefore decreasing the volume similarity. Indeed, Iglesias et al. (2016) found an average of 4.5% difference in absolute volume similarity in their test-retest condition. In terms of Dice, all other methods were at a disadvantage to FS Long for this metric due to the interpolation step required to realign the scans. This effect was mitigated through the utilisation of a fuzzy Dice coefficient in all other methods. However, despite the disadvantage, LASHiS shows comparable Dice overlaps in the test-retest condition to FS Long, except for the smaller subfields (e.g., CA2/3). One additional potential source of bias is the various anatomical definitions of hippocampus subfields proposed by both FS Long and ASHS/LASHiS. We attempted to correct for differences in anatomical priors of subfields by combining subfields that were present in both ASHS and Freesurfer. However, it should be noted that the definition of the anatomical priors by the anatomists/raters of the respective atlas packages would have a strong influence on segmentation results, and in particular, Dice overlaps. For example, utilising an atlas package that intrinsically defines smaller subfields would result in thinner subfields, resulting in lower Dice scores. However, previous work by Xie et al. (2018) has shown that atlas composition largely does not affect the segmentation quality of 7 T images using ASHS. The influence of label priors of Freesurfer has not been examined, so we therefore cannot exclude the possibility that the Freesurfer method is biased negatively by the initial labelling of the atlas. Coupled with the results of volume similarity, we can assert that LASHiS is a reliable method for longitudinal hippocampus subfield segmentation.

### 4.2 Experiment Two, Bayesian longitudinal Linear Mixed Effects modelling

Many evaluation strategies employ manual segmentations (e.g., Berron et al., 2017) to provide a gold standard for evaluation of any segmentation strategy. However, manual hippocampus subfield segmentation is time and labour intensive, taking up to eight hours initially, and two hours after five months of training (Wisse et al., 2016) and is prone to inter- and intra-rater variability (Boccardi et al., 2011; Hsu et al., 2002; Mulder et al., 2014). We explored the usefulness of LASHiS in the examination of the variance ratio in our longitudinal Bayesian LME modelling experiment. Higher variance ratios that are characterised by both lower residual variability and larger between-subjects variability are beneficial for longitudinal cohort studies. In the TOMCAT dataset, we found the highest variance ratios in LASHiS, underscoring the usefulness of our approach in maximising between subject differences. We note outliers in variance ratios in LASHiS and FS Long, which are driven by results in CA2-3, and SUB, respectively. For LASHiS, high residual variability was found for right CA2-3, driving this outlier.

For the ADNI data, only one subfield (right DG) had a significantly higher variance ratio in LASHiS than FS Long, with all other subfields having overlapping 95% CIs. Our result in the ADNI dataset suggests that using any longitudinal pipeline is advantageous for examining hippocampal subfield volumetry. Optimised longitudinal pipelines such as Fs Long and LASHiS are important for analysing data. However, the atlas package should be the determining factor when selecting which pipeline to utilize. For LASHiS in ADNI, we made use of the Penn Memory Center ASHS atlas, which has been labelled on older aged participants with similar contrast and acquisition parameters to the data acquired in ADNI. We therefore suggest that at 3 T, the choice of atlas composition and labelling scheme should inform the user on segmentation method choice.

We want to note here, that LASHiS is potentially negatively biased by limitations in subfield selection. All Freesurfer schemes combine CA2 & CA3 estimates in their algorithms. In calculating our subfield estimates, we summed CA2 and CA3 volumes offline, potentially biasing our estimates of residual variability. We note a similar residual variability outlier in the ASHS Xs scheme in the left CA2-3 combined subfield regions. Volume estimates of CA2 and CA3 regions were generally reported less precisely than other subfields, as measured by the low test-retest statistics and low within-subject variability in the LME experiment. Previous research (Dalton et al., 2017; Pipitone et al., 2014; Wisse et al., 2016; Yushkevich et al., 2015) has repeatedly shown discrepancies in reporting these subfield boundaries *in vivo*. This is largely due to their small size and the reliance on heuristic geometric rules for segmenting CA2/3 subfields on *in vivo* MRI, rather than visible contrast differences in the scan. Thus, inter- and intra-rater reliability are often low for these subfields (Xie et al., 2018). Our automatically derived subfield estimates are likely influenced by discrepancies in the manual labels that inform segmentations. Notably, FS Long also suffers from a low variance ratio in CA2-3, suggesting either i) a homogeneous participant pool leading to low between-subject variability, ii) large, unexpected differences in time points in these subfields, or iii) a combination of these.

### 4.4 Benefits and advantages of LASHiS

Both LASHiS and the FS Long scheme segment hippocampus subfields and derive volume estimates from MRI images. However, only the T1w scan of an individual is processed through the longitudinal stream of ‘recon-all’ before longitudinal processing of hippocampus subfields, potentially explaining the FS Long results compared to LASHiS. Our design utilises multi-contrast information from MRI scans and importantly allows for information that can *only* be captured by multi-contrast MRI (i.e., the subfields of the hippocampus) to be included in the labelling.

LASHiS derives its power from its ability to decrease random errors in the labelling procedure, and through increasing the likelihood for correct labelling to occur when the SST is created. This implicitly increases SNR and sharpness of the SST compared to the individual time points through the template building procedure (Shaw et al., 2019). Our inclusion of Diet LASHiS highlights the contributions of the JLF step from the simple labelling of the SST, which may be subject to both random and systematic errors. In LASHiS, these random errors may be mitigated in part due to the bootstrapping of JLF from the individual time point to the SST. It is possible that this step decreases the likelihood of random errors in the labelling scheme because of JLFs ability to vote on labels that fit best to the SST. Therefore, random variance caused by mislabelling at any individual time point may be ameliorated by the JLF step. In turn, this is the likely cause for the low residual variability found in LASHiS in Experiment Two. Our inclusion of JLF labelling using *automatically* generated labels is a novel consideration in the field of hippocampus subfield segmentation and relies on the assumption that automatically generated subfield labels are considered accurate.

We included a computationally less expensive and faster approach to multi-contrast hippocampus subfield segmentation, namely Diet LASHiS. This method performs all steps save for the initial cross-sectional segmentations and the bootstrapping of these segmentations to the SST using JLF. Diet LASHiS performed well in the volume similarity portion of Experiment 1, and in Experiment 2 in comparison to the other methods examined, though to a lesser degree than LASHiS in the TOMCAT dataset. In the ADNI dataset, Diet LASHiS performs worse than both LASHiS and Fs Long, suggesting the reverse normalisation to SST approach may not be suitable for smaller subfield estimations at 3 T (as the small CA2&3 regions largely drove this result). It is possible that the SST creation step in Diet LASHiS is not optimal for 3 T data, the larger voxel size may incur a larger partial volume effect when inverting the deformation field. As the steps taken to complete LASHiS and Diet LASHiS are the same except for the additional JLF bootstrapping method, we conclude that the increased sensitivity and robustness in the LASHiS scheme was due to the JLF step. Indeed, despite the disadvantage of potentially increasing systematic errors with the JLF bootstrapping step, it is evident that these systematic errors are largely overcome in the initial cross-sectional labelling of the hippocampus subfield with ASHS Xs.

Processing time for LASHiS depends largely on compute infrastructure, T2w image size, and the number of time points. Our testing on three time points with high-resolution (0.3 mm^3^) T2w images ran in the order of 24 hours on a single CPU core, and 4 hours on 8 cores and 64 GB memory without parallelisation. Many steps, including the initial cross-sectional segmentations and SST creation, can be run in parallel using job scheduling software (PBS, Sun Grid Engine, Slurm, etc.) and parallelised across cores, decreasing the time required by orders of magnitude commensurate with the number of cores employed. Diet LASHiS is estimated to decrease compute time by approximately 20%, as neither the cross-sectional, nor the JLF steps are required. ASHS Xs takes between 1-2 hours on a single core, while FS Xs takes approximately 40 minutes after 24-48 hours of pre-processing on a single core (https://surfer.nmr.mgh.harvard.edu/fswiki/ReconAllRunTimes). Fs Long takes approximately 60 minutes on a single core after 24 hours of cross-sectional processing per time point, and further creation of an SST. On eight cores with 64 GB of memory and with parallelisation, our average run time for the entire Freesurfer longitudinal pipeline was 20 hours per participant. The great advantage of LASHiS is the flexibility of computational processing options for each step, allowing for scalable processing of larger datasets.

Our incorporation of a Bayesian approach to the widely used longitudinal LME method for examining differences in method performance aids in discrimination of subtle differences between participants with small variability (as in the present study). This technique allowed us to simultaneously examine small differences between participants, while also capturing longitudinal within-subject changes; both of which are especially important in examining clinical subpopulations and other low-*n* studies, where small longitudinal changes need to be captured precisely.

### 4.5 Limitations

The design of our pipeline decreases random variability in any session due to the SST registration and JLF scheme. A limitation of our scheme is that label errors (i.e., systematic errors) in subjects will propagate to the SST, despite the sophisticated JLF algorithm employed that does not independently compute similarity weights between the pairs while voting (H. Wang et al., 2013). Therefore, it is important to note that LASHiS is never free from labelling errors that occur in all image segmentation pipelines. These systematic errors can be avoided through quality assurance of scans and labels at the cross-sectional level (i.e., before the JLF bootstrapping step), which is essential in any volumetric labelling scheme, regardless.

We here report a small healthy cohort of young adults with no known psychological or neurological disorders scanned at 7 T. We assumed no difference between time point two and three, and very small differences between time point one and two due to the age and health of the participants. We concede this limitation in our interpretation of test-retest analyses. The Bayesian nature of our longitudinal LME modelling accounts for small sample sizes (Sorensen et al., 2016), and the results of Experiment One and Two should therefore not be affected by our small sample size in the TOMCAT dataset. We also included a larger dataset from the ADNI consortium, though were unable to conduct test-retest statistics on this sample due to data availability. Future work could improve upon the methods we used for the ADNI dataset, including building and manually labelling an atlas with the specific intensity profile/MR characteristics of ADNI data in order to improve segmentation priors for LASHiS.

Our test-retest statistics in the 7 T TOMCAT dataset show that LASHiS has improved metrics compared to other longitudinal methods, with obvious differences to previous work reporting the same methods (Iglesias et al., 2016), where Dice overlaps were considerably higher overall for the Freesurfer methods. We note this limitation of having such a small sample size in the present study, which was the likely reason for the higher variability in the Dice overlap scores in the Freesurfer method. However, as LASHiS shows a consistent improvement compared to all other methods, we are confident LASHiS is a robust and reliable method for longitudinal multi-contrast hippocampus subfield segmentation.

### 4.6 Conclusions

Here, we present a technique for automatically and robustly segmenting hippocampus subfield volumes using UHF multi-contrast MRI in healthy subjects. We found that LASHiS shows marked improvements at 7 T across several relevant measures, such as Dice similarity and volume similarity coefficients for test-retest reliability, and Bayesian LME modelling, compared to other methods used for cross-sectional and longitudinal hippocampus segmentation. Results from the 3 T ADNI dataset highlight the importance of utilising longitudinal pipelines for hippocampus volumetry, with the user determining pipeline choice by the atlas priors. LASHiS utilises multi-contrast information and joint-label fusion, which better captures hippocampus subfield tissue characteristics and decreases random errors in the labelling procedure.

## Supporting information

Supplementary Figures for LASHiS

## Acknowledgements

The authors acknowledge the facilities and scientific and technical assistance of the National Imaging Facility, a National Collaborative Research Infrastructure Strategy (NCRIS) capability, at the Centre for Advanced Imaging, The University of Queensland. MB acknowledges funding from Australian Research Council Future Fellowship grant FT140100865. This research was undertaken with the assistance of resources and services from the Queensland Cyber Infrastructure Foundation (QCIF). The authors would like to acknowledge Siemens Healthineers for the support of our project through the provision of the WIP sequence used to acquire the tse_UHF_WIP729C (variant: tse2d1_9) and MP2RAGE sequence (WIP 900) data in this publication. Data collection and sharing for this project was funded by the Alzheimer’s Disease Neuroimaging Initiative (ADNI) (National Institutes of Health Grant U01 AG024904) and DOD ADNI (Department of Defense award number W81XWH-12-2-0012). ADNI is funded by the National Institute on Aging, the National Institute of Biomedical Imaging and Bioengineering, and through generous contributions from the following: AbbVie, Alzheimer’s Association; Alzheimer’s Drug Discovery Foundation; Araclon Biotech; BioClinica, Inc.; Biogen; Bristol-Myers Squibb Company; CereSpir, Inc.; Cogstate; Eisai Inc.; Elan Pharmaceuticals, Inc.; Eli Lilly and Company; EuroImmun; F. Hoffmann-La Roche Ltd and its affiliated company Genentech, Inc.; Fujirebio; GE Healthcare; IXICO Ltd.; Janssen Alzheimer Immunotherapy Research & Development, LLC.; Johnson & Johnson Pharmaceutical Research & Development LLC.; Lumosity; Lundbeck; Merck & Co., Inc.; Meso Scale Diagnostics, LLC.; NeuroRx Research; Neurotrack Technologies; Novartis Pharmaceuticals Corporation; Pfizer Inc.; Piramal Imaging; Servier; Takeda Pharmaceutical Company; and Transition Therapeutics. The Canadian Institutes of Health Research is providing funds to support ADNI clinical sites in Canada. Private sector contributions are facilitated by the Foundation for the National Institutes of Health (www.fnih.org). The grantee organization is the Northern California Institute for Research and Education, and the study is coordinated by the Alzheimer’s Therapeutic Research Institute at the University of Southern California. ADNI data are disseminated by the Laboratory for Neuro Imaging at the University of Southern California.

