## Supplementary Figures for LASHiS for "Longitudinal Automatic Segmentation of Hippocampal Subfields (LASHiS) using Multi-Contrast MRI"

#### Supplementary Materials One: Experiment One

We conducted a series of Bayesian paired-sample t-tests in order to test absolute volume differences between the second and third time point. Supplementary Figure 1 and 2 show the differences between methods and subfields for volume similarity in the test-retest experiment.

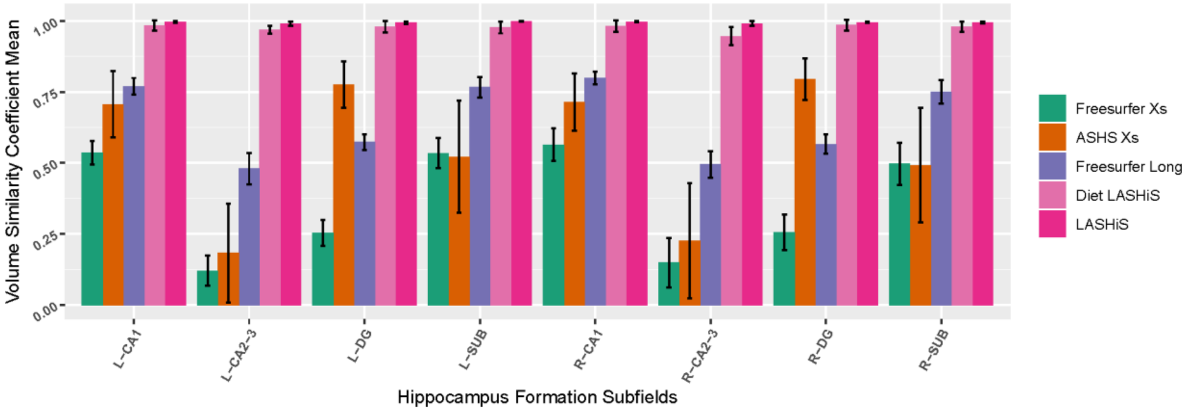


*Supplementary Figure 1. Bar plots of volume similarity coefficients of each hippocampus subfield (x axis) between time point two and time point three of the TOMCAT dataset. Different methods are denoted by different colours. Error bars represent standard deviations.*

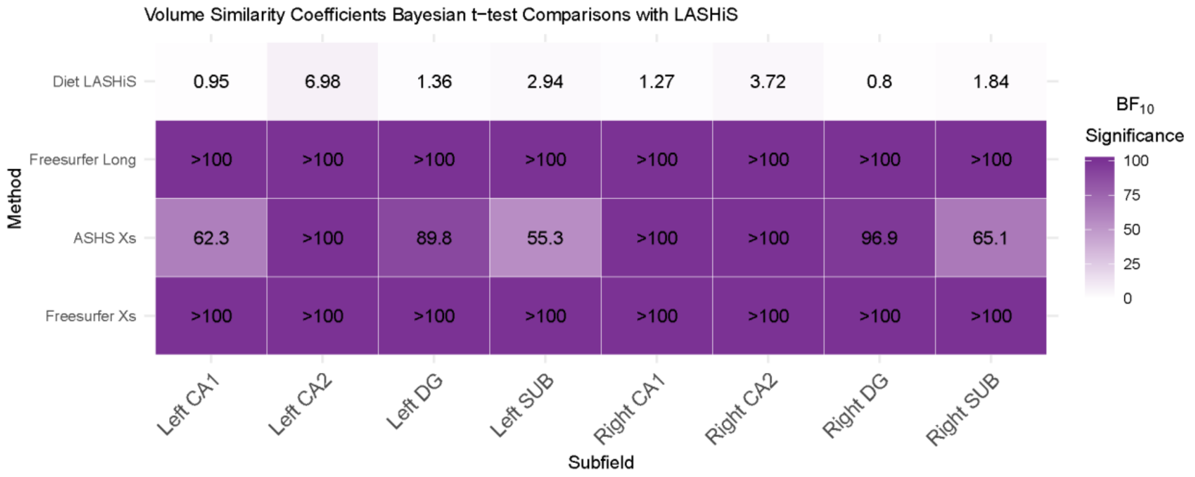


*Supplementary Figure 2. Matrix of significance values for volume similarity comparisons between LASHiS and the other four methods (y-axis). Each subfield was compared to LASHiS for volume similarity using Bayesian t-tests, and the $BF_{10}$ values*

*are reported here. $BF_{10}$ > 3 was taken as substantial evidence for the alternative hypothesis, with $BF_{10}$ > 10 taken as strong evidence, and $BF_{10}$ greater than 100 were considered decisive. $BF_{10}$ values between 1 and 3 were considered anecdotal evidence for the alternative hypothesis. In contrast, $BF_{10}$ < 0.33 was considered as substantial evidence for the null, with $BF_{10}$ between 0.33 and 1 providing anecdotal evidence for the null hypothesis in accordance with (Lee & Wagenmakers, 2013).*

We next conducted Bayesian paired-sample *t*-tests for Dice overlaps between the segmentation labels in the second and third time point. Supplementary Figure 2 shows Dice overlap values of each subfield for each method. Note, that Dice scores for LASHiS, Diet LASHiS, Freesurfer Xs and ASHS Xs are negatively affected by the resampling needed to compute the registrations between the two time points, which is not present in the FS Long method.

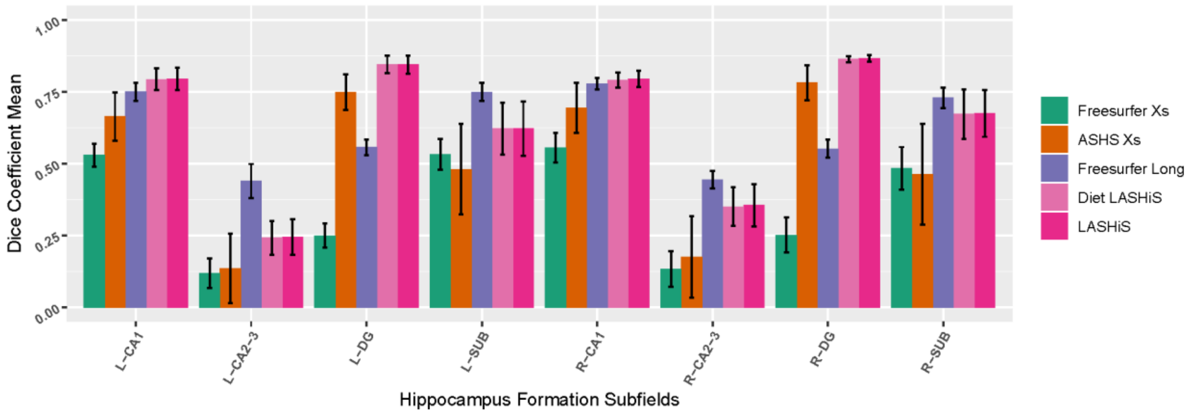


*Supplementary Figure 3*. *Bar Plots of Dice coefficients of each hippocampus subfield (x-axis) between time point two and time point three of the TOMCAT dataset. Each method is shown in a different colour. A value of 1 represents perfect overlap between time points, and 0 represents no overlap. Freesurfer Xs, ASHS Xs, Diet LASHiS and LASHiS all require resampling to a common space before overlap calculation of Dice overlaps. Higher scores between time points denote higher subfield overlap between test-retest time points. Error bars represent standard deviations.*

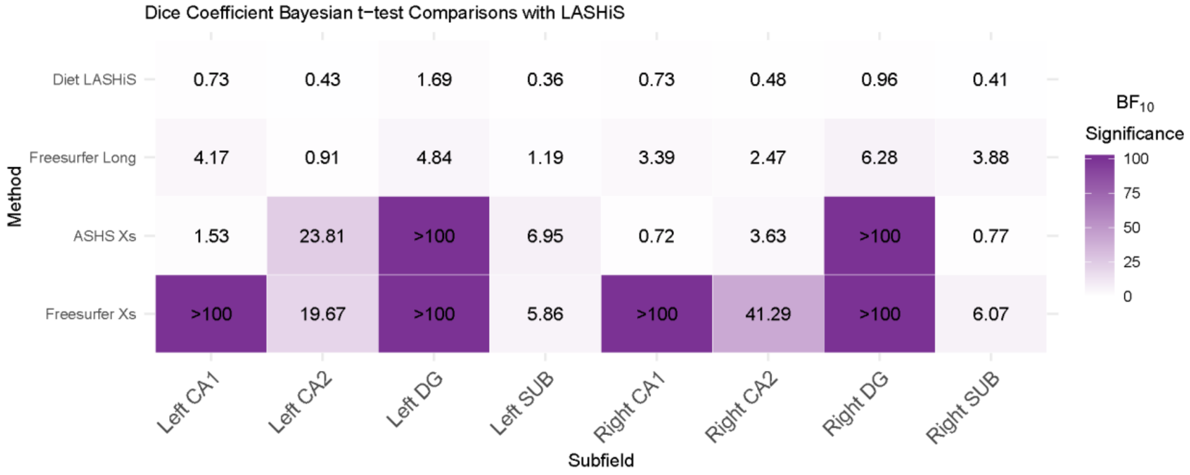


*Supplementary Figure 4. Matrix of significance values for Dice coefficient comparisons between LASHiS and the other four methods (y-axis). Each subfield was compared to LASHiS for Dice using Bayesian t-tests, and the $BF_{10}$ values are reported here. $BF_{10}$ > 3 was taken as substantial evidence for the alternative hypothesis, with $BF_{10}$ > 10 taken as strong evidence, and $BF_{10}$ greater than 100 were considered decisive. $BF_{10}$ values between 1 and 3 were considered anecdotal evidence for the alternative hypothesis. In contrast, $BF_{10}$ < 0.33 was considered as substantial evidence for the null, with $BF_{10}$ between 0.33 and 1 providing anecdotal evidence for the null hypothesis in accordance with (Lee & Wagenmakers, 2013).*

#### Supplementary Materials Two: Experiment Two

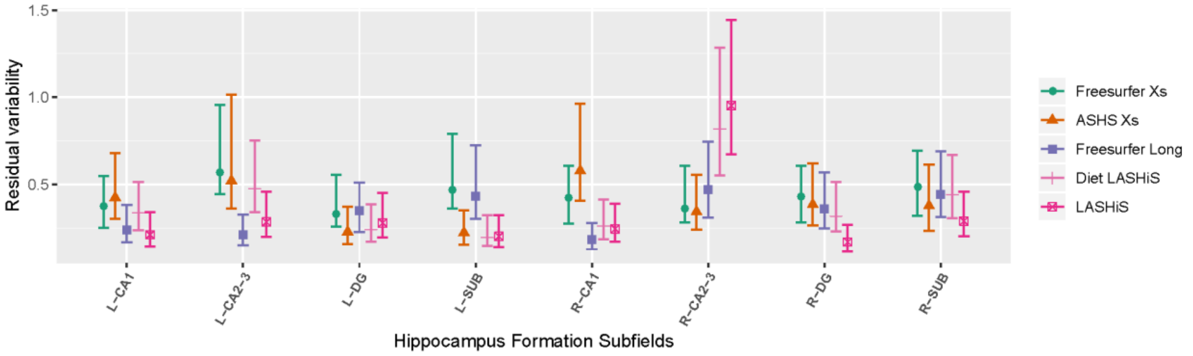


*Supplementary Figure 5. Residual (within-subjects) variability for each subfield (x-axis), for each method (coloured lines). Lower values represent decreased variability of the method between time point two and time point three. Shapes represent the mean residual variability, with lines denoting the 95% confidence intervals for each method.*

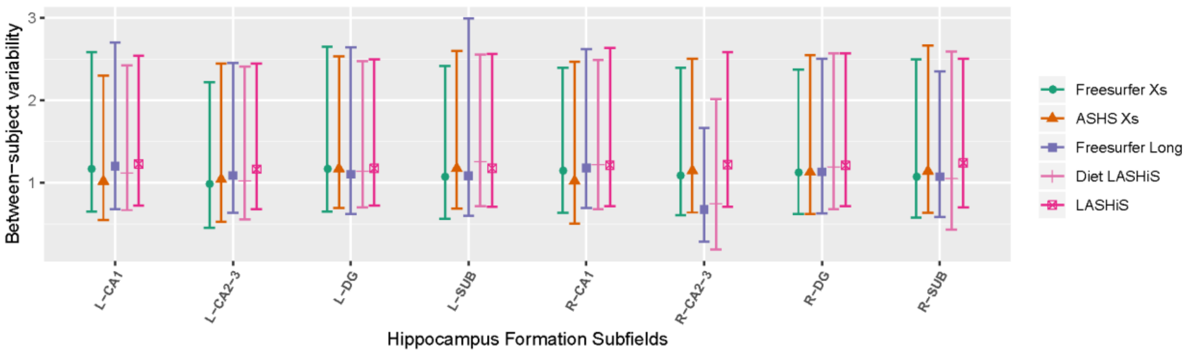


*Supplementary Figure 6. Between-subjects variability for each subfield (x-axis), for each method (coloured lines). Higher values represent increased variability of the method between time point two and time point three and a larger difference between participants, indicating a pipeline's ability to discriminate between participants more successfully. Shapes represent the mean between-subjects variability, with lines denoting the 95% confidence intervals for each method.*

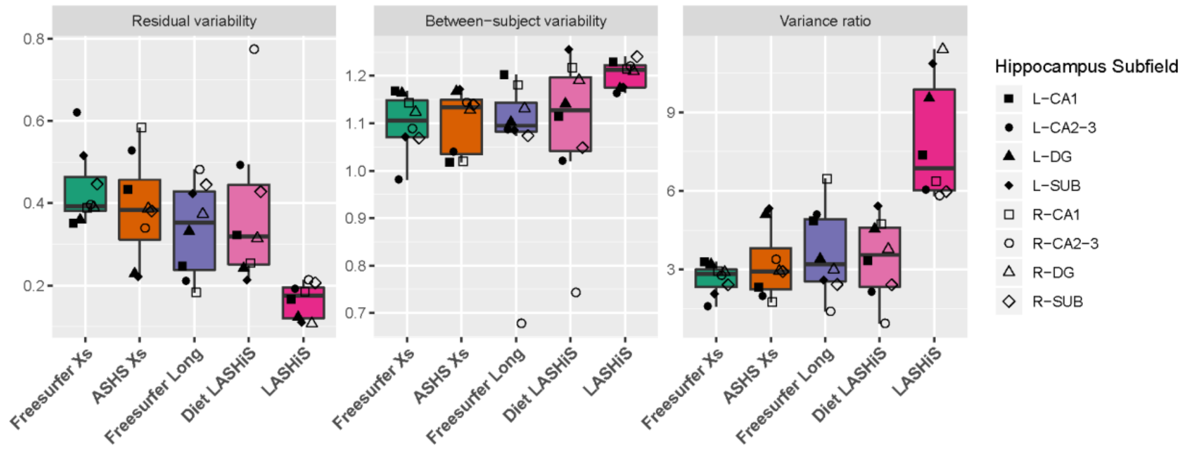


*Supplementary Figure 7. Box plots of (from left to right) residual variability, between-subject variability, and the variance ratio between the two variabilities of each method for the three time points for the TOMCAT (7 T) dataset. Error bars denote the 95% confidence intervals. Shapes with black and white fill represent individual left, and right hippocampus subfields, respectively. Lower residual, and higher between-subject variabilities are preferred for longitudinal pipelines. The variance ratio is a summary statistic of the two variability values, with higher values indicating improved discrimination between within- and between-subject variance.*

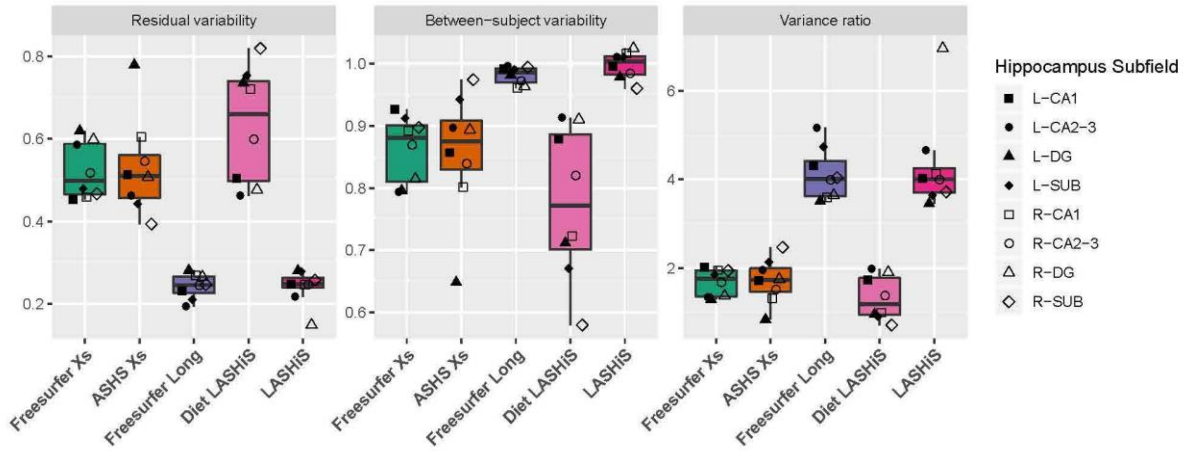


*Supplementary Figure 8. Box plots of (from left to right) residual variability, between-subject variability, and the variance ratio between the two variabilities of each method for the three time points for the ADNI (3 T) dataset. Error bars denote the 95% confidence intervals. Shapes with black and white fill represent individual left, and right hippocampus subfields, respectively. Lower residual, and higher between-subject variabilities are preferred for longitudinal pipelines. The variance ratio is a summary*

*statistic of the two variability values, with higher values indicating improved discrimination between within- and between-subject variance.*

**Supplementary Materials Three: LASHiS code example.**

*From https://github.com/thomshaw92/LASHiS*

### Requirements:

Requires ANTs >v2.3.0 https://github.com/ANTsX/ANTs/

Requires ASHS https://sites.google.com/site/hipposubfields/home

LASHiS performs a longitudinal estimation of hippoocampus subfields. The following steps are performed:

1. Run Cross-sectional ASHS on all time points
2. Create a single-subject template (SST) from all the data, then cross-sectionally run the SST through ASHS.
3. Using the Cross-sectional inputs as priors, label the hippocampi of the SST.
4. Segmentation results are reverse normalised to the individual time point.

### Environment Variables:

ASHS_ROOT Path to the ASHS root directory

ANTSPATH Path to the ANTs root directory

### Misc Notes:

LASHiS was loosely adapted from the ANTs Longitudinal Cortical Thickness pipeline https://github.com/ANTsX/ANTs/.The ASHS_TSE image slice direction should be z. In other words, the dimension of ASHS_TSE image should be 400x400x30 or something like that, not 400x30x400

### Usage:

```
/path/to/LASHiS.sh -a atlas selection for ashs \
      <OPTARGS> \
      -o outputPrefix \
      \${anatomicalImages[@]} \
```

### Required arguments:

```
-o:  Output prefix                                    The following subdirectory and
images are created for the single
                                                     subject template
                                                       *
\${OUTPUT_PREFIX}SingleSubjectTemplate/
                                                       *
\${OUTPUT_PREFIX}SingleSubjectTemplate/T_template*.nii.gz
 -a: Atlas selection                                Full path for the atlas you would
like to use for the Cross-sectional
                                                     labelling of ASHS and the SST.
Can be made in ASHS_train
 anatomical images                                 Set of multimodal (T1w or gradient
echo, followed by T2w FSE/TSE input)
                                                     data. Data must be in the format
specified by ASHS & ordered as follows:
                                                       \${time1_T1w}   \${time1_T2w}
\\
                                                       \${time2_T1w}   \${time2_T2w}
...
                                                       .
                                                       .
                                                       .
                                                        \${timeN_T1w}  \${timeN_T2w}
...
```

### Optional arguments:

```
-c:  control type                                   Control for parallel computation
for ANTs steps (JLF,SST creation)  (default 0):
                                                       0 = run serially
                                                       1 = SGE qsub
                                                       2  =  use  PEXEC  (localhost)
(remember to define cores in -j)
                                                       3 = Apple XGrid
                                                       4 = PBS qsub
                                                       5 = SLURM

 -d:  OPTS                                             Pass in additional options to
SGE's qsub for ASHS. Requires -c 1

 -e:  ASHS file                                        ProConfiguration file. If not
passed, uses $ASHS_ROOT/bin/ashs_config.sh
 -f:  Diet LASHiS                                    Diet LASHiS (reverse normalise
the SST only) then exit.

 -g:  denoise anatomical images                    Denoise anatomical images (default
= 0).
 -j:  number of cpu cores                                Number of cpu cores to use
locally for pexec option (default 2; requires "-c 2")
```

```
 -n:  N4 Bias Correction                     If yes, Bias correct the input
images before template creation.
                                            0 = No
                                            1 = Yes

 -b:  keep temporary files                   Keep warps, ASHS Xs files, etc
(default = 0).
```
